# *Burkholderia pseudomallei* clinical isolates are highly susceptible *in vitro* to cefiderocol, a novel siderophore cephalosporin

**DOI:** 10.1101/2020.03.26.009134

**Authors:** Delaney Burnard, Gemma Robertson, Andrew Henderson, Caitlin Falconer, Michelle Bauer-Leo, Kyra Cottrell, Ian Gassiep, Robert Norton, David L. Paterson, Patrick N. A. Harris

## Abstract

Cefiderocol is a novel cephalosporin designed to treat multidrug resistant Gram-negative infections. By forming a chelated complex with ferric iron, cefiderocol is transported into the periplasmic space via bacterial iron transport systems and primarily binds to penicillin-binding protein 3 (PBP3) to inhibit peptidoglycan synthesis. This mode of action results in cefiderocol having greater *in vitro* activity against many Gram-negative bacilli than currently used carbapenems, β-lactam/β-lactamase inhibitor combinations, and cephalosporins. Thus, we investigated the *in vitro* activity of cefiderocol (S-649266) against a total of 271 clinical isolates of *Burkholderia pseudomallei* from Australia. The collection was comprised of primary isolates (92.3%) and subsequent isolates (7.7%). Minimum inhibitory concentrations (MIC) of cefiderocol ranged from ≤0.03 to 32 mg/L, where the MIC_90_ was 1 mg/L and 16 mg/L for primary and subsequent isolates, respectively. Based upon non-species specific (Gram-negative bacilli) clinical breakpoints for cefiderocol (MIC ≤ 4 mg/L), twelve isolates (4.4%) would be classified as non-susceptible. Further testing for co-resistance to meropenem, ceftazidime, trimethoprim-sulfamethoxazole, amoxicillin-clavulanate and doxycycline was performed on a subset of isolates with elevated cefiderocol MICs (≥2 mg/L, 4.8%) and 84.6% of these isolates exhibited resistance to at least one of these antimicrobials. Cefiderocol was found to be highly active *in vitro* against *B. pseudomallei* primary clinical isolates. This novel compound shows great potential for the treatment of melioidosis in endemic countries and should be explored further.

## Introduction

Pathogenic multidrug-resistant (MDR) and carbapenem-non-susceptible Gram-negative bacilli (GNB) have expanded significantly worldwide^1,2^, diminishing appropriate antimicrobial therapy options^3,4^. Cefiderocol (formerly S-649266, Shionogi & Co. Ltd., Osaka, Japan) inhibits peptidoglycan synthesis, and has been described as almost ubiquitously stable against β-lactamases, resulting in greater efficacy than carbapenems, currently available β-lactam/β-lactamase inhibitor combinations and cephalosporins^5–8^.

Cefiderocol recently received FDA approval for the treatment of complicated urinary tract infections^9^, on the basis of non-inferiority against imipenem/cilastatin^10^. A trial versus meropenem for nosocomial pneumonia has also been completed^5,11^ with efficacy demonstrated against Gram-negative pathogens such as Enterobacterales, *Pseudomonas aeruginosa* and *Acinetobacter baumannii*^12,13^. Provisional CLSI *in vitro* breakpoints have been set at ≤4 mg/L (S) and ≥16 mg/L (R) respectively for these pathogens^12^. Furthermore, cefiderocol has shown promising efficacy against *Burkholderia pseudomallei* and *B. mallei* in limited sample sizes (*n*= 30), in the recommended Iron-depleted cation-adjusted Mueller-Hinton broth (ID-CAMHB) (MIC_90_ 0.25, 4 mg/L), respectively^14^.

*B. pseudomallei* is endemic to tropical and sub-tropical regions, including south-east Asia and northern Australia and suspected for Africa and South America^15–17^. The bacteria is known to cause melioidosis, where pneumonia, sepsis, neurological disease and visceral abscesses are commonly described clinical presentations^15,18^. Treatment for *B. pseudomallei* infections require an “intensive” 2-8-week intravenous treatment with an antimicrobial such as a carbapenem or cephalosporin, then followed by and orally administered “eradication” treatment for 3-6 months with an antimicrobial such as trimethoprim-sulfamethoxazole^19^. Infections frequently require intensive care admission and are associated with significant morbidity^20^. This is, in part due to the intrinsic antimicrobial resistance (AMR) of *B. pseudomallei*, as well as acquired resistance to antimicrobials such as tetracyclines, β-lactam/β-lactamase inhibitors and rarely carbapenems^21–23^. The mortality rate for patients presenting with melioidosis ranges from 10-30% in Burma, Singapore, Thailand and Vietnam^20^, while in Australia it remains approximately 10%^24^. Recent research has focused on new or repurposed compounds in order to provide improved treatment options for such infections, particularly given the potential of *B. pseudomallei* to be genetically manipulated or used as an agent of biologic warfare^25,26^. Therefore, with the need for new treatment options for *B. pseudomallei* and other GNB, and promising efficacy of cefiderocol as a treatment option, the aim of this study was to assess the *in vitro* activity of cefiderocol against a large sample size of clinical *B. pseudomallei* isolates from endemic regions of Australia.

## Methods

### Ethics

Ethical approval for this study was granted by the Forensic and Scientific Services Human Ethics Committee (HREC/17/QFSS/12). Biosafety approvals for this study were granted by the Institutional Biosafety Committee UQCCR (IBC/210B/SOM/2017). This project was performed under the study number: S-649266-EF-312-N.

### Isolates

Clinical *B. pseudomallei* isolates were prospectively collected from patients admitted to Queensland Health hospitals, Australia, over a period from 1999 to 2018. A small number of isolates were referred from external laboratories. These isolates were retrieved from −80°C storage from three microbiology laboratories in Queensland (Forensic and Scientific Services, Coopers Plains; Pathology Queensland, Townsville and Central laboratories). Isolates were transferred to the University of Queensland Centre for Clinical Research (UQCCR) and stored at stored at −80°C prior to testing. Demographic and clinical information for the isolates and patients was retrieved from the laboratory information system (Auslab; PJA Solutions).

### Disc Diffusion

Disc diffusion susceptibility testing was performed using 30 µg cefiderocol discs provided by Mast Group Ltd. (Bootle, UK). Discs were stored at 4°C prior to use. Isolates were sub-cultured from storage on 5% horse blood agar (Micromedia, Victoria, Australia) for 18-24h at 37°C prior to preparation of a 0.5 McFarland solution in 0.9% sterile saline. Mueller-Hinton agar plates (Micromedia, Victoria, Australia) were inoculated with test isolates using the Kirby-Bauer method and incubated for 16-20h at 37°C. Control strains *E. coli* ATCC 25922 and *P. aeruginosa* ATCC 27853 were included with each run.

### Broth Microdilution

Broth microdilution (BMD) was performed using 96-well plates provided by Shionogi & Co., Ltd. (Osaka, Japan) and prepared by International Health Management Associates (IHMA; Schaumburg, USA). Iron-depleted cation-adjusted Mueller-Hinton broth (ID-CAMHB) was used according to CLSI and manufacturer recommendations^27^. Plates were stored at −20°C and thawed for one hour prior to use. Isolates were sub-cultured from storage on to 5% horse blood agar (Micromedia, Victoria, Australia) for 18-24h at 37°C prior to preparation of a 0.5 McFarland solution in 0.9% sterile saline for a final inoculum of 5 × 10^5^. A maximum of three plates were inoculated and then sealed during each run, owing to biosecurity restrictions. Plates were incubated at 37°C for 16 to 20h prior to reading. One positive growth control well and one negative control well were included in each plate. Quality control of cefiderocol was performed using *E. coli* ATCC 25922 and *P. aeruginosa* ATCC 27853 strains, with only those runs that passed QC and with appropriate growth in the positive control included for analysis^28^. CLSI provides susceptible (≤4 mg/L) and resistant (≥16 mg/L) interpretative criteria for Enterobacterales and some non-fermenters such as *Pseudomonas aeruginosa, Acinetobacter baumannii* and *Stenotrophomonas maltophilia* (but not *B. pseudomallei*)^12^. As such, these breakpoints were applied to isolates in this study.

Further antimicrobial susceptibility testing was performed on isolates with elevated cefiderocol MICs, for meropenem, ceftazidime, doxycycline, amoxicillin-clavulanic acid and trimethoprim-sulfamethoxazole. BMD was performed in house where 96 well plates were prepared using the Tecan D300e Digital Dispenser (HP Inc. CA, USA.) Standard inoculum (5 × 10^5^ CFU), incubation in ambient air at 37°C and end point reading at 24 hours were performed. Quality control was performed in duplicate for each batch of plates made including *E. coli* ATCC 25922, *E. coli* ATCC 35218 and *S. aureus* ATCC 29213 to ensure correct dilutions of each antimicrobial^28^. The range of meropenem, ceftazidime, doxycycline, amoxicillin-clavulanic acid and trimethoprim-sulfamethoxazole tested was 0.06-16, 0.06-16, 0.12-16, 0.12/2-32/2 and 0.06/1.19-32/608 mg/L, respectively. MICs were interpreted according to CLSI breakpoints^29^.

## Results

### Isolates

A total of 271 isolates from 250 individuals were included in this study (Table 1a & b). Multiple isolates from 15 patients were identified and separated into primary (n=250) and subsequent (n=21) isolate test groups. Subsequent isolates comprised of duplicate specimens, specimens collected from an alternate anatomical site or isolates collected weeks, months or years apart. The antimicrobial treatment of patients is unknown. From the 250 individuals, *B. pseudomallei* was predominantly isolated from males and blood, lung, skin and soft tissue specimens (Table 1a & b). All isolates were from Queensland with the exception of one from New South Wales, two from the Northern Territory, two from South Australia and one from Tasmania. International isolates included: one each from England and Germany, two from New Zealand and two from Papua New Guinea.

**Table 1a.** Demographic and clinical data for primary *B. pseudomallei* isolates.

| *Individuals* |  |  |
| --- | --- | --- |
|  | *n* patients | 250 |
|  | Age range | 6-90 years |
|  | Gender | 26.4% F, 73.6% M |
| *Isolate collection site* |  |  |
|  | Blood | 135 |
|  | Lung | 31 |
|  | Skin and soft tissue | 27 |
|  | Urine | 14 |
|  | Sputum | 11 |
|  | Unknown | 6 |
|  | Bone/joint | 5 |
|  | Liver | 2 |
|  | Synovial fluid | 2 |
|  | Prostate | 1 |
|  | Brain | 1 |
|  | Lymph node | 1 |
|  | Gastrointestinal tract | 1 |
|  | Gut | 1 |
|  | Pleural fluid | 1 |
|  | Peritoneal fluid | 1 |
| *Isolate total* | *n* | 250 |

**Table 1b.** Demographic and clinical data for secondary and subsequent *B. pseudomallei* isolates.

| *Individuals* |  |  |
| --- | --- | --- |
|  | *n* patients | 15 |
|  | Age range | 10-74 years |
|  | Gender | 53.3% F, 46.6% M |
| *Isolate collection site* |  |  |
|  | Lung | 8 |
|  | Blood | 5 |
|  | Skin and soft tissue | 4 |
|  | Unknown | 2 |
|  | Bone/joint | 1 |
| *Isolate total* | *n* | 21 |

### Disc diffusion

All 271 isolates were subject to disc diffusion testing. Zone diameters ranged from 14-46 mm and 11-43 mm for primary and subsequent isolates, respectively (Table 2). Two isolates C137 (38 mm) and T62 (41 mm) demonstrated inner growth on disc diffusion testing (12 mm and 17 mm respectively), indicating a heterogeneous population or expression.

**Table 2.** *In vitro* minimum inhibitory concentrations (mg/L) and disc diffusion range (mm) for both primary and subsequent *B. pseudomallei* isolates against cefiderocol.

| Isolates | Cefiderocol MIC (mg/L) |  |  |  |  |  |  |  |  |  |  |  | Disc Zone |  |  |
| --- | --- | --- | --- | --- | --- | --- | --- | --- | --- | --- | --- | --- | --- | --- | --- |
|  | ≤0.03 | 0.06 | 0.12 | 0.25 | 0.5 | 1 | 2 | 4 | 8 | 16 | ≥32 | Range | MIC<sub>50</sub> | MIC<sub>90</sub> | (mm) |
| Primary<br>(n=250) | 72 | 93 | 0 | 13 | 32 | 35 | 0 | 3 | 1 | 1 | 0 | ≤0.03-8 | 0.06 | 1 | 14-46 |
| Subsequent<br>(n=21) | 6 | 4 | 0 | 1 | 2 | 0 | 1 | 1 | 4 | 1 | 1 | ≤0.03-32 | 0.25 | 16 | 11-43 |

### Broth Microdilution

Cefiderocol BMD was performed on all 271 isolates. For primary isolates the MIC range, MIC_50_, and MIC_90_ were ≤0.03-8, 0.06 and 1 mg/L, while the corresponding values for subsequent isolates were ≤0.03-32, 0.25 and 16 mg/L (Table 2, Figure 1). Significant differences between zone diameter and MIC were only observed for MIC values ≥8 mg/L, in both isolate groups (Figure 2).

**Figure 1.**
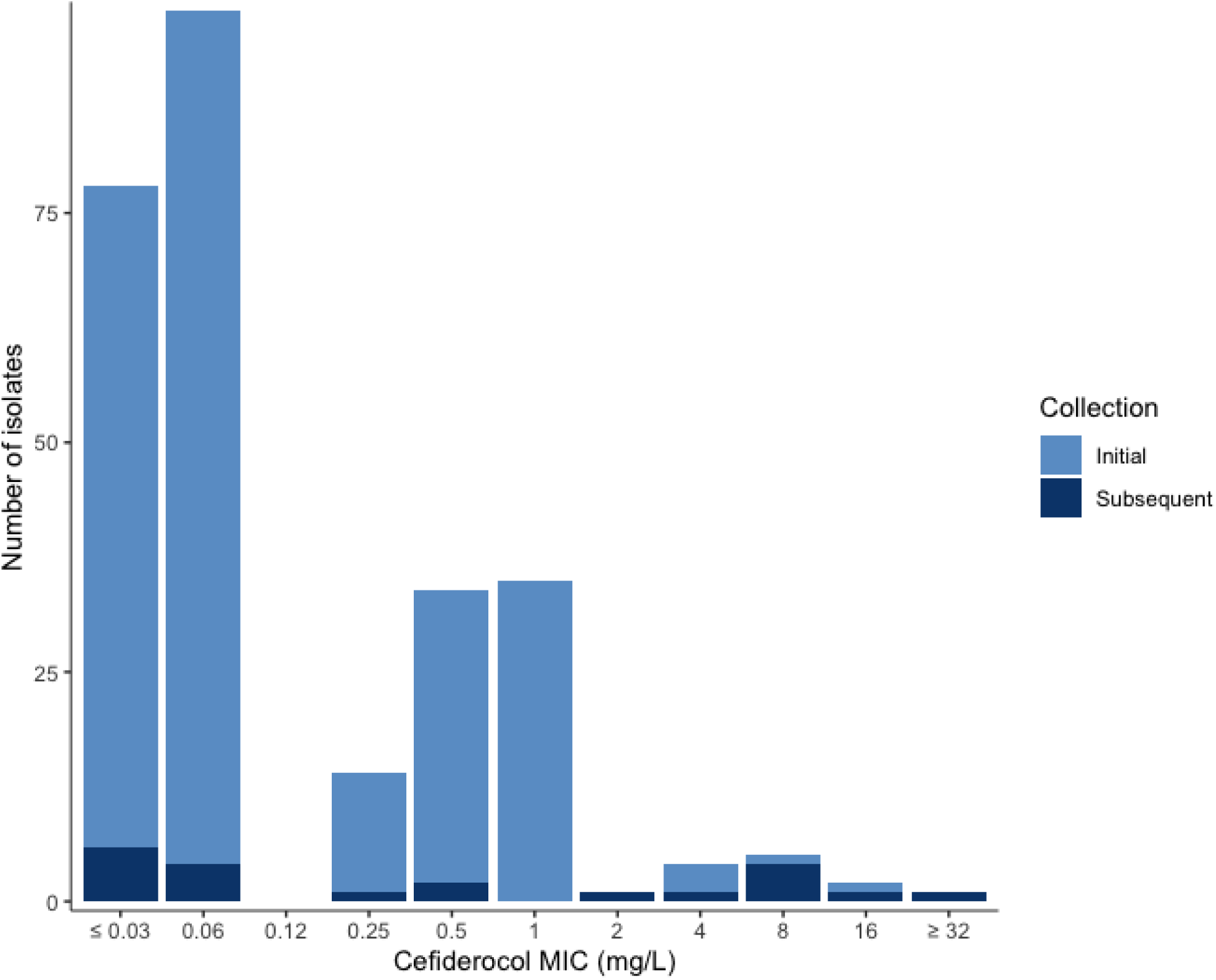
Stacked histogram of cefiderocol minimum inhibitory concentrations (mg/L) for 271 clinical *Burkholderia pseudomallei* isolates. Primary isolates collected are shown in light blue, while subsequent isolates are shown in dark blue.

**Figure 2.**
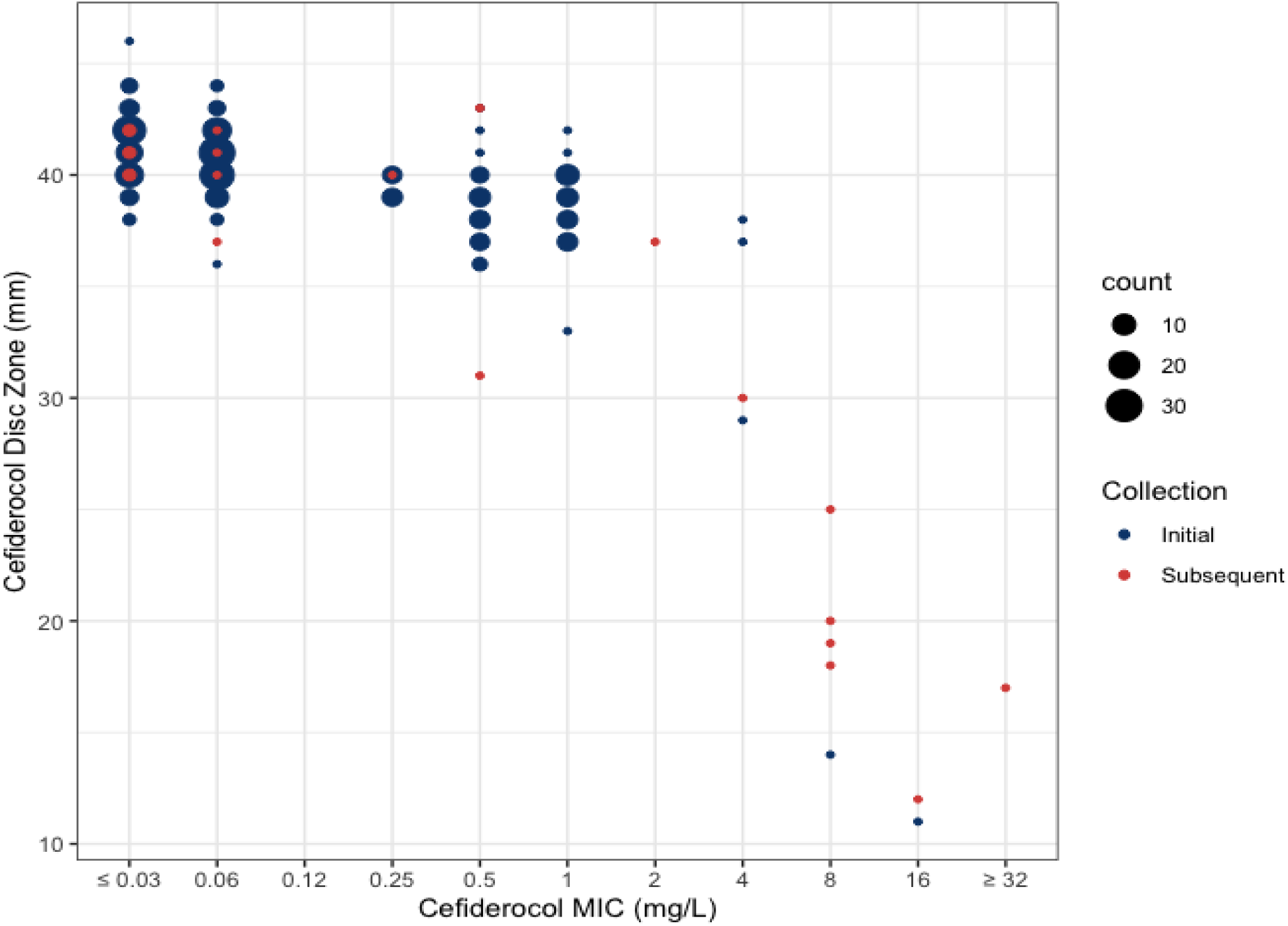
Scatterplot of cefiderocol disc zone diameters (mm) vs cefiderocol minimum inhibitory concentrations (mg/L) for 271 clinical *Burkholderia pseudomallei* isolates. Primary isolates collected are shown in blue, while subsequent isolates are shown in red.

### Isolates expressing elevated cefiderocol minimum inhibitory concentrations

According to CLSI breakpoints for cefiderocol against other GNB, 8 isolates (2.9%) would be categorised as non-susceptible (≥8 mg/L). However, our data displayed a tri-modal distribution (Figure 1) where isolates with MICs ≥2 mg/L represented the 2.5% of isolates with the highest MICs. This included four isolates at 4 mg/L, one at 2 mg/L and 8 isolates with MICs ≥8 mg/mL, resulting in 13 isolates (4.8%) being considered to have elevated MICs (i.e ≥2 mg/L) in this study. The 13 isolates (4.8%) with elevated cefiderocol MICs were comprised of 46.1% primary and 53.9% subsequent isolates, predominantly collected from the lung (53.9%) and individuals with serious underlying co-morbidities (92.3%) (Table 3). Co-resistance was examined via BMD between cefiderocol and meropenem, ceftazidime, amoxicillin clavulanic acid, doxycycline, and trimethoprim-sulfamethoxazole; with 11 of the 13 isolates (84.6%) also resistant to amoxicillin-clavulanic acid. Resistance to the other antimicrobials tested was observed, yet inconsistently (Table 3, Figure 3).

**Table 3.** Minimum inhibitory concentrations of *B. pseudomallei* isolates with elevated cefiderocol (CEF) concentrations against meropenem (MEM), ceftazidime (CAZ), amoxicillin-clavulanic (AMZ) acid, trimethoprim-sulfamethoxazole (SXT) and doxycycline (DOX). Non-susceptible and resistant categories are denoted with NS or R, respectively. Isolates from individuals with underlying co-morbidities are noted.

| *B. pseudomallei* isolate | Site collected | Underlying co-morbidity | Clinically relevant antimicrobials (mg/L) |  |  |  |  |  |
| --- | --- | --- | --- | --- | --- | --- | --- | --- |
| Primary |  |  | CEF | MEM | CAZ | AMC | SXT | DOX |
| C4 | Lung | Yes | 8 | □ 16 R | 2 | □ 32 R | 0.12/2.28 | 2 |
| T17 | Blood | Unknown | 4 | 2 | ≥ 16 NS | □ 32 R | 0.25/4.75 | ≥ 16 R |
| T18 | Blood | Yes | 2 | 2 | 4 | □ 32 R | 0.12/2.28 | 1 |
| C50 | Lung | Yes | 4 | 4 | ≥ 16 NS | □ 32 R | 4/76 R | 4 |
| T63 | Sputum | Yes | 4 | 1 | 4 | □ 32 R | 0.12/2.28 | 8 |
| C194 | Blood | Yes | 16 | □ 16 R | 2 | □ 32 R | 0.12/2.28 | 2 |
| Subsequent |  |  | CEF | MEM | CAZ | AMC | SXT | DOX |
| C5 | Lung | Yes | 8 | □ 16 R | □ 16 NS | □ 32 R | 16/304 R | □ 16 R |
| C6 | Lung | Yes | 8 | 1 | □ 16 NS | 4 | 0.12/2.28 | 8 |
| C7 | Lung | Yes | 8 | 4 | 1 | 4 | 0.12/2.28 | 8 |
| C15 | Lung | Yes | 8 | 1 | 2 | 32 R | 16/304 R | 16 R |
| C52 | Lung | Yes | 4 | 4 | □ 16 NS | □ 32 R | 4/76 R | 4 |
| T62R | SSTI | Unknown | 32 | 1 | 4 | □ 32 R | 0.25/4.75 | 2 |
| C137R | SSTI | Yes | 16 | 1 | 2 | □ 32 R | 0.5/9.5 | 2 |

**Figure 3.**
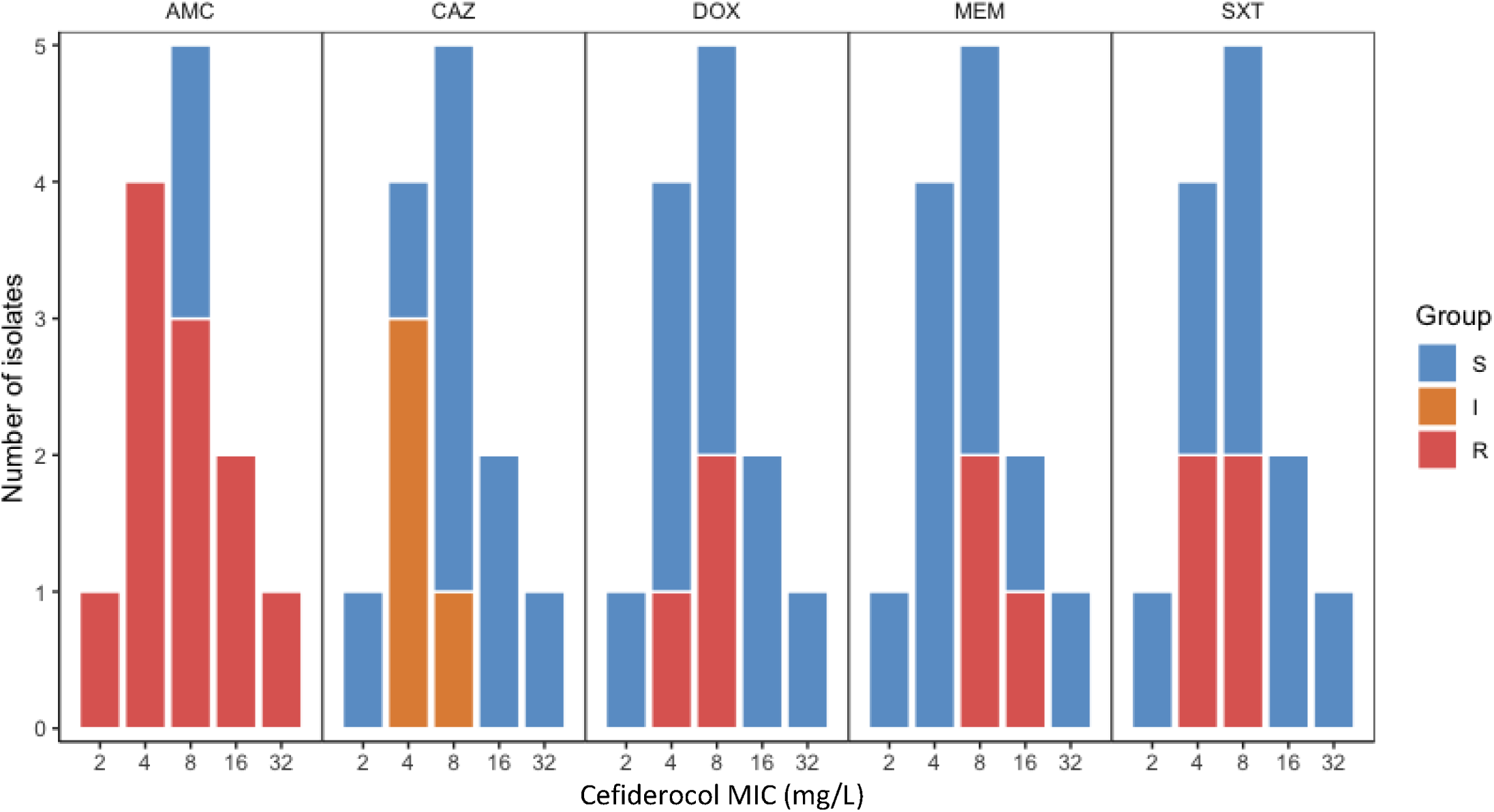
Stacked histogram of cefiderocol minimum inhibitory concentrations (mg/L) against categorised amoxicillin-clavulanic acid (AMC), ceftazidime (CAZ), doxycycline (DOX), meropenem (MEM) and trimethoprim-sulfamethoxazole (SXT) minimum inhibitory concentrations (mg/L) for *Burkholderia pseudomallei* isolates with cefiderocol MIC values of 2 mg/L or greater.

## Discussion

*B. pseudomallei* infections pose a significant mortality risk to patients. Furthermore, treatment can be especially burdensome in terms of the need for prolonged intravenous and oral antibiotic therapy. Due to the virulence of the organism, and its predilection for immune compromised hosts, patients with melioidosis frequently present with severe disease requiring intensive care admission. While uncommon, emergent resistance may also occur to compromise treatment. A novel compound such as cefiderocol with such low MICs, could be used as an empirical therapy to improve clinical outcomes of *B. pseudomallei* in endemic regions, where multi-drug resistant Gram-negative bacilli are circulating. An approach such as this, may result in improved clinical responses during empirical therapy and increased bacterial killing in hard to treat infections like those in the central nervous system, bone, joint and lymphatic system.

### Disc diffusion

The relationship between cefiderocol disc diffusion zone diameter and BMD MIC values was observed at concentrations ≥8 mg/L of cefiderocol. As there are no zone diameters specified for cefiderocol disc diffusion against *B. pseudomallei* very major errors, major errors and minor errors were unable to be calculated. Zone diameter was able to differentiate resistant (≥8 mg/L) isolates consistently at less than 25mm (Figure 2). This relationship was previously observed in over 1,300 Gram-negative bacteria where resistant isolates (≥8 mg/L) were consistently differentiated at 20 mm or less^30^. These relationships were present in species such as Enterobacteriaceae, *P. aeruginosa* and *A. baumannii*, however *S. maltophilia* isolates (≥2 mg/L) were differentiated at 15 mm or less ^30^. Better discrimination among susceptible isolates may be seen with a lower disk mass, but only 30 µg disks are currently available commercially.

### Broth microdilution

In this study isolates with MIC values ≥2 mg/L were considered to have elevated cefiderocol MICs, yet few were identified in this collection (*n*=13). This finding is expected, with previous *in vitro* studies identifying very few resistant isolates for several Gram-negative bacilli species, for example only 24/753 isolates showed MICs ≥8 mg/L from a global dataset^31^, 54/8954 isolates had MICs ≥4 mg/L^13^ and no resistant isolates were identified among 189 isolates from Greece^32^. *In vitro* MIC_90_ values of the 250 primary *B. pseudomallei* isolates was 1 mg/L in this study, higher than the previously described MIC_90_ of 0.25 mg/L obtained from a pilot study of 30 *B. pseudomallei* isolates^14^. The increase in MIC may be influenced by the increase in sample size, however larger samples sizes have not substantially increased the MIC_90_ of cefiderocol in other pathogens such as *P. aeruginosa, A. baumannii, K. pneumoniae* and *E. coli*^13,31,32^. Moreover, sister species *B. mallei* has an MIC_90_ of 4 mg/L from 30 isolates, significantly higher than that observed in *B. pseudomallei*^14^, whereas isolates from the closely related *B. cepacia* complex also exhibit a consistently low MIC_90_ of 0.016 and 0.12 mg/L, despite small sample sizes in ID-CAMHB media^5,13^.

### Isolates with elevated cefiderocol minimum inhibitory concentrations

As no patient had received cefiderocol previously, we expect the MIC increase is likely a reflection of including isolates from complex patients with persistent or relapsing infection. Subsequent isolates exhibited an exceptionally high MIC_90_ of 16 mg/L with 84.6% of these isolates originating from patients with underlying conditions. Six isolates C50 and 52 (4 mg/L) and C4, 5, 6, and 7 (8 mg/L) were derived from the sputum of a patient with cystic fibrosis or prolonged chronic *B. pseudomallei* lung colonisation. These isolates have been exposed to significant and prolonged treatment with antimicrobials, representing and influencing 46.1% of the isolates with elevated cefiderocol MICs observed. Furthermore, five isolates C15, C137R, C194, T18 and T63 were derived from patients with immunosuppression following organ transplantation or poorly controlled diabetes, representing and influencing 38.5% of *B. pseudomallei* isolates with elevated cefiderocol MICs. Clinical data was not available for the remaining two isolates. Our results predict elevated cefiderocol MICs will more than likely be encountered in individuals with prolonged or extensive antimicrobial exposure, in conjunction with persistent or relapsing *B. pseudomallei* infections.

Five clinically relevant antimicrobials were selected for further BMD testing against *B. pseudomallei* isolates with elevated cefiderocol MICs, to assess the presence of co-resistance. Interestingly, 84.6% of the 13 isolates exhibited resistance to amoxicillin-clavulanic acid, followed by 38.5% with non-susceptibility to ceftazidime, 30.8% with trimethoprim-sulfamethoxazole resistance and 23% with meropenem and doxycycline resistance, respectively. These rates of resistance are much higher than those reported from Australia (0.0-4.0%) in previous studies^33–35^. Subsequent isolate C5 exhibited a multidrug resistant profile with resistance to meropenem, amoxicillin-clavulanic acid, trimethoprim-sulfamethoxazole, doxycycline, non-susceptibility to ceftazidime and a cefiderocol MIC of 8 mg/L. Previous studies have highlighted that cefiderocol activity is not generally impacted by resistance to other antimicrobials^5–7,13,32^. In this study, amoxicillin-clavulanic acid resistance does not appear to impact cefiderocol or ceftazidime MICs. As these isolates are from individuals with known antimicrobial exposure and significant co-morbidities, it is less likely that elevated cefiderocol MICs reflect co-resistance, and is more likely a result of exposure and adaptation.

### Prospective role of cefiderocol against B. pseudomallei infections

Cefiderocol has great potential as antimicrobial therapy for multi-drug resistant *B. pseudomallei* infections. This is supported by the findings of this study, in combination with the efficacy of cefiderocol being unaffected by resistance to other antimicrobials *in vitro*^32^, the antimicrobial being well tolerated by patients^5,8^ and found to be safe in clinical trials to date^8^. However, this study also suggests that cefiderocol may lose efficacy *in vitro* against *B. pseudomallei* isolates derived from individuals with significant underlying co-morbidities and prolonged antibiotic exposure. Nevertheless, further *in vitro* testing, followed by *in vivo* trials of cefiderocol against *B. pseudomallei* is certainly validated.

## Conclusion

Cefiderocol demonstrates a high degree of activity *in vitro* against 271 clinical isolates of *B. pseudomallei* from Australia. The MIC_50_ and MIC_90_ are 0.06 and 1 mg/L, respectively. Elevated cefiderocol MICs were infrequently demonstrated in this collection (4.8%) based on non-species-specific breakpoints and is most commonly associated with significant underlying co-morbidities. Cefiderocol shows promise as an intravenous agent for the management of acute melidoidosis based upon *in vitro* susceptibility testing, particularly in regions where carbapenem-resistant Gram-negative organisms and *B. pseudomallei* co-circulate. Further investigation into the role of cefiderocol as a treatment for melidoidosis and likely mechanisms of resistance would be of great value.

## Acknowledgements

The authors would like to acknowledge the ongoing and financial support of Shionogi & Co., LTD., under the stand-alone service agreement. Additionally, the authors would like to acknowledge the efforts of Pathology Queensland collaborators: Haakon Bergh and Tracey Shepherd during this study.

